# An Analysis of Silk Density in Spider Webs

**DOI:** 10.1101/2025.01.26.634972

**Authors:** Fangyuan Lin, Kathryn M. Nagel, Seewoo Lee, Jason Jiang, Grant Yang, Patrick Chang, Samuel Li, Norman Sheu

## Abstract

This study explores the structural complexity of spider webs through information-theoretic and harmonicity-based frameworks to quantify spatial patterns in silk density across different web regions and reveal the underlying resource allocation strategies. Currently, there is no normalized approach for describing web structure and complexity, particularly for sheet-webs, and this methodology allows for non-destructive scanning and quantification of web characteristics. By analyzing the entropy of silk density distributions, a single scalar that captures the heterogeneity of material investment across the entire web, we observed that the entropy values follow a normal distribution with a mean of 1.24 ± 0.22 bits when using 10 quantization levels. In the second part of the paper, by measuring the harmonicity of the silk density, we reveal that the silk density at a given point can be inferred from its neighbors, with an average harmonicity value of 0.0039 ± 0.0017 (fraction of total points in point cloud data). The harmonic behavior is notable for its maximum principle, suggesting that the strongest parts of the web appear at the boundaries, aligning with existing knowledge of spider web construction. These findings provide a new technique for quantifying web-building strategies and offer new insights into spider behavior and evolution.

## 1 Introduction

Spiders are exceptional biological engineers, using self-manufactured silk to design and build webs many times larger than their bodies [1]. As a material, spider silk is stronger and more elastic than almost any other known natural and man-made material, while simultaneously being biodegradable and biocompatible [2, 3, 4, 5]. Spiders use silk for a variety of purposes, including prey capture and wrapping, encasing egg sacks, during courtship and copulation, and perhaps most visibly: building webs [6]. A spider’s web serves as a tool for prey capture, an extension of the senses, and a retreat from predators [6, 7, 8, 9, 10]. Based on the most up-to-date phylogenetic analysis, orb webs are the most ancient type of prey capture web, evolving more than 165 Mya [11, 12, 13, 14]. Whether the web evolved once or multiple times is still under debate, but regardless of the origin, many extant families abandoned orb webs later in their evolution, adopting cursorial lifestyles or a variety of other web structures, including sheet webs, cob-webs, and funnel webs [6, 15]

The linyphioids group (Linyphiidae and Pimoidae) constitute about 10 percent of all spider species worldwide, with more than 5000 named species [16]. Despite linyphioid diversity and abundance, web architecture and mechanics are poorly studied. Linyphiids generally build clearly suspended horizontal sheet webs that are sparsely attached to the surrounding substrate; however, variations in this design are prevalent in most genera [17]. Sheet web structures are generally described as an unordered meshwork of silk fibers varying in thickness and strength, with some threads bearing viscid droplets [7, 18]. However, building a web is an energetically costly process, depleting glucoproteins and significantly increasing the metabolic rate [19, 20]. It is expected that over evolutionary time, selection would favor economical webs that maximize function with the least amount of silk [21, 22]. However, quantifying the effectiveness and economy of webs is a difficult process with no universally accepted metric, although mathematical modeling has precedence for cob and orb web species [23, 24, 25, 26].

Another important aspect in spider web complexity is the spatial correlations in silk density [26]. By examining how the density of silk in one region of the web relates to the density in neighboring regions, we can infer patterns and predict the distribution of silk throughout the web. Understanding how silk densities vary across the 3D surface of a sheet web can potentially provide further insight into spider web building behavior and identify areas of the web used for different purposes, for example, prey capture compared to predator retreat [6, 17].

In this paper, we develop a method to quantify sheet web complexity by measuring Shannon entropy and harmonicity for the sheet webs of three species of linyphiid common to Northern California, *Neriene digna, Neriene litigiosa*, and *Microlinyphia dana*. Using these metrics, we are able to discern new features of sheet webs and deepen our understanding of spider web-building behavior.

The remainder of the paper is organized as follows. In Sections 2 and 3, we provide background information on the mathematical principles and experimental methodology, respectively. Section 4 discusses entropy calculations and the corresponding statistical analyses. Section 5 discusses the harmonicity of silk density and the analysis of local spatial correlations across the webs. Finally, Section 6 concludes the paper and suggests directions for future research.

## 2 What are Entropy and Harmonicity?

### 2.1 Mathematical Framework for Entropy

#### 2.1.1 Motivation for Entropy

Quantities that capture macroscopic properties of a system are valuable since they enable meaningful analysis without requiring a detailed account of the system’s microscopic structure. For example, temperature measures the thermal energy of a substance without reference to the motion of individual atoms or molecules. Silk density in a spider web is directly proportional to the spider’s material investment and energetic cost of construction, but there is currently no standard metric for quantifying this information aside from simple summary figures such as total silk mass [27, 1].

By quantizing local silk-mass measurements into a discrete distribution and computing their Shannon entropy, we obtain a single scalar that captures the heterogeneity of that investment across the entire web. This macroscopic value allows us to quantify global organizational features of the web while abstracting away the intricate and often irregular details of individual thread placement, and allows for comparison between webs of different sizes and structures. Crucially, our entropy-based measure is also insensitive to the absolute scale or the particular choice of segmentation, which facilitates robust comparisons across webs of different sizes, shapes, and species.

This methodology shows that in all scanned webs, 98-99% of the space is empty and have zero density value, leaving us with an apparent sparsity of data to analyze. However, this is due to the implicit nature of spider webs, which are by definition minute amounts of silk occupying otherwise empty space. By removing the zero density values from our analysis, we are focusing our analysis and observations on only the biological structure, rather than excluding or failing to describe any part of the web itself.

#### 2.1.2 Mathematical Framework for Entropy Computation

Generally, Shannon entropy measures the amount of information needed to describe the spread of data across all possible values [28]. By examining the silk density distribution across a web, we can assign an entropy value to each web. A low entropy value would indicate that the silk density across the web is primarily concentrated at one value, while a high entropy value would indicate a web with a full range of density values spread uniformly at random. The lowest possible entropy value is zero bit, and the highest is log_2_ (10) ≈ 3.3219 bits when quantizing with 10 density levels.

We develop a mathematical framework for computing the entropy of silk density distributions. The silk density distribution of a web can be determined by subdividing the three-dimensional space where the web is constructed by a cubic grid and then quantizing the silk density measurements into discrete levels. It should be noted that this framework demonstrates a convergence property: as we refine the subdivisions of the spatial domain, the resulting entropy becomes asymptotically independent of how the region is subdivided.

Following Shannon’s definition of entropy, we compute entropy of the density of a web as follows: Divide the whole box with measurements given by height, width, and depth into *N = N*_height_ × *N*_width_ × *N*_depth_ smaller boxes of size 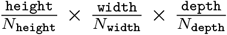. For 0 ⩽ *i N*_height,_ 0 ⩽ *j* < *N*_width,_ 0 ⩽ *k* < *N*_depth_ let *n*_*i,j,k*_ be the number of points in the corresponding small box *B*_*i,j,k*_. Then the local density of a box is

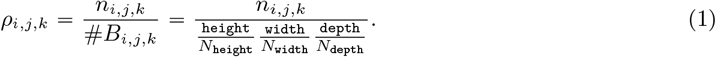

We group these densities into *L* different levels (bins), and compute the entropy of the corresponding distribution. To do this, we first compute *ρ*_max_, the greatest density among all boxes. Then we sub-divide the range from 0 to *ρ*_max_ into *L* equal intervals. Each density can then be categorized into its corresponding interval. A box *B*_*i,j,k*_ is assigned to level 0 if and only if *n*_*i,j,k*_ 0, i.e. it contains no point.

##### Definition 2.1

(Entropy of a web’s silk density). Define the *entropy* of a web *W* ‘s silk density as

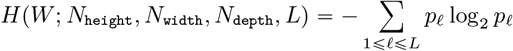

where

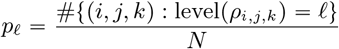

are the probability densities for each level *ℓ* ∈ {1, 2, …, *L*}. Observe that empty boxes (level 0) do not enter into the entropy sum since they are not part of the web.

Note that our definition of entropy depends on the choice of the subdivisions *N*_height_, *N*_width_, *N*_depth_ and the number of levels *L*. However, we remark that in theory, the entropy converges as we take finer subdivisions.

##### Theorem 2.2

(Convergence of Silk Density Entropy with Respect to Subdivision). Assuming there exists a measurable underlying field *ρ* of silk density over the box, as the subdivisions become finer (i.e., *N*_height_, *N*_width_, *N*_depth_ → ∞), the entropy *H*(*W* ; *N*_height_, *N*_width_, *N*_depth_, *L*) converges to a finite limit *H*(*W* ; *L*) ∈ ℝ _⩽0_.

*Proof*. Since Shannon’s entropy is continuous with respect to the variational distance *L*^1^-distance), it is enough to show that the empirical distribution of silk density levels converges in *L*^1^ to the true distribution of the density levels as the number of subdivisions *N = N*_height_ × *N*_width_ × *N*_depth_ goes to infinity. The average silk density *ρ*_*i,j,k*_ of the *ijk*-th subdivided box *B*_*i,j,k*_ can be written as an integral 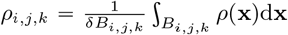 (here *δB*_*i,j,k*_ is a volume of *B*_*i,j,k*_), and Lebesgue’s differentiation theorem implies that *ρ*_*i,j,k*_ converges to *ρ*(**x**) as *N* → ∞. Since silk density functions (both empirical and true) are measurable, one can use Lebesgue differentiation theorem to show that *ρ*_*i,j,k*_ converges to *ρ* (**x**) as *δB*_*i,j,k*_ → 0, i.e. *N* → ∞. The level function is also measurable, hence the indicator function for each level of empirical density distribution converges to the indicator function of the true density distribution by dominated convergence theorem. By summing over all levels, we can conclude the *L*^1^ convergence.

Figure 1 shows the convergence of entropy as the number of subdivisions grows, which empirically verifies the claim of Theorem 2.2.

**Figure 1.**
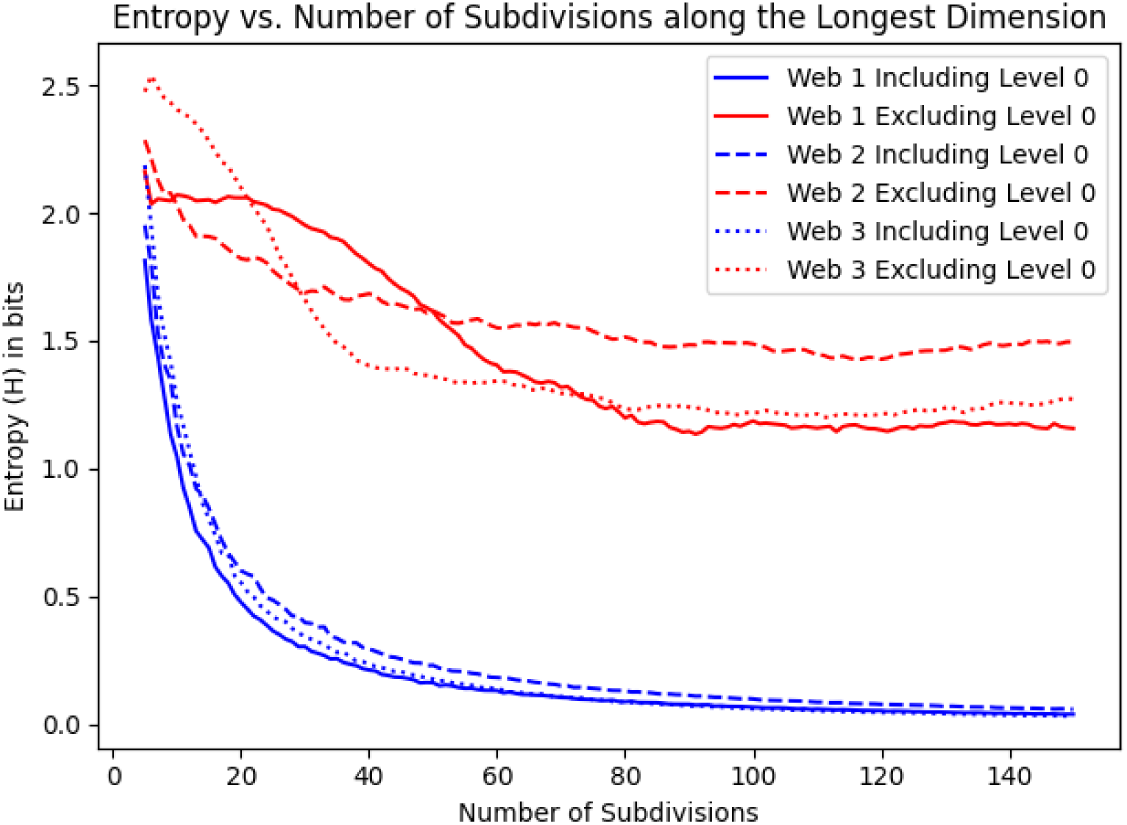
Convergence of entropy with respect to number of subdivisions along each dimension for three different webs. The spider that constructed web 1 and web 3 is a randomly selected juvenile female *M. dana* The spider that constructed web 2 is a randomly selected male *M. dana*.

#### 2.1.3 Choice of the Number of Quantization Levels

For the following experiments, we choose *L =* 10 for the number of levels. To justify our choice density-levels, we both provide a theoretical bound and show that, in practice, entropy values increase linearly if we use *L =* 100 levels instead and therefore, increasing the number of quantization levels from 10 to 100 does not yield additional information (Figure 2). The theorem below demonstrates that raising the number of levels from *L* to *mL* increases the entropy by at most log_2_ *m*. As a result, we find *L =* 10 sufficiently fine to capture all relevant heterogeneity while remaining computationally efficient.

**Figure 2.**
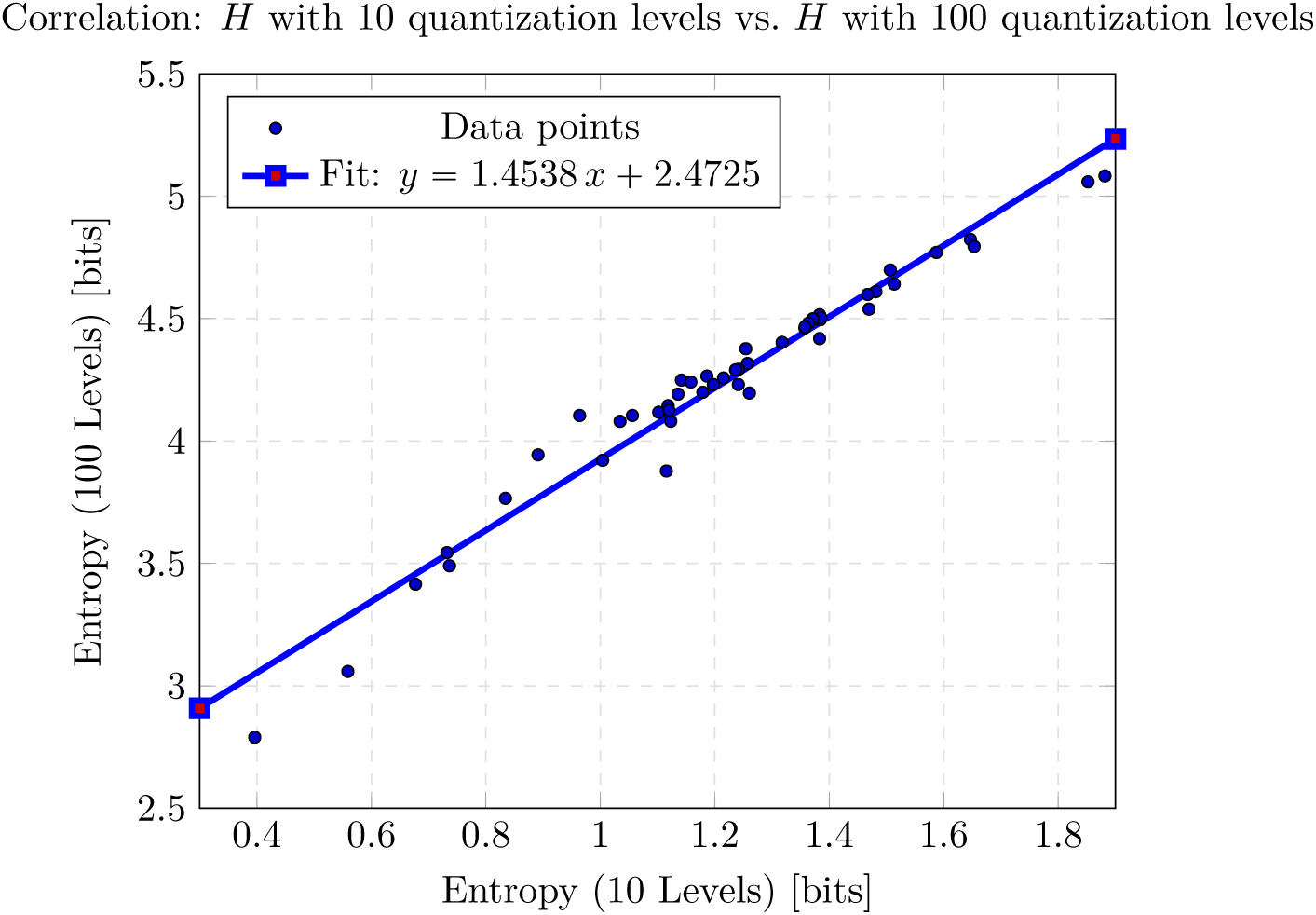
Scatter of *H*_100_ versus *H*_10_ with fitted regression line *Ĥ*_100_ = 1.4538 *H*_10_ + 2.4725. The model explains 95.9% of the variance (*R*^2^ = 0.9589), with a slope highly significant (*t* = 32.76, *p* < 10^−32^) and a residual standard error of 0.09195 bits (*n* = 48).

##### Theorem 2.3.

Fix *N*_height_, *N*_width_, *N*_depth_ and let *H*(*W* ; *L*) = *H*(*W, N*_height_, *N*_depth_, *N*_width_, *L*). For an integer *m* ⩽ 1, we have

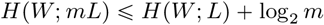

*Proof*. Let *n*_*ℓ*_ (resp. 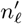) be the number of samples (local densities) of level *ℓ* with maximum level *L* (resp. *mL*). Then we have 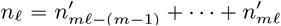 (the bin corresponds to the level *ℓ* is partitioned into *m*-bins with new levels *m*_*ℓ*_ − (*m* − 1), *mℓ* − (*m* − 2), …, *m*_*ℓ*_). By Jensen’s inequality (applied to *f* (*x*) = − *x* log_2_ *x*), we have

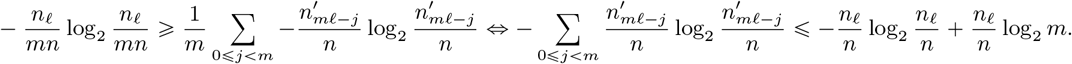

Then we get

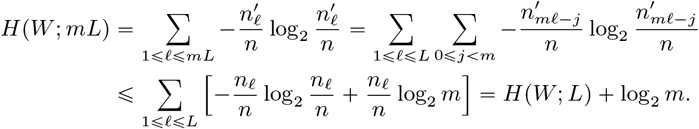

Figure 2 is a scatter plot showing the correlation between the distribution of entropy with *L* = 10 and *L* = 100 levels, which shows that the distributions are highly positively correlated.

#### 2.1.4 Visualization of Low and High Entropy Webs

The following figures present heat maps of the silk density distributions for two samples: one with a relatively low entropy value for the silk density distribution and the other with a relatively high entropy value.

In the low-entropy web (Figure 3, left), silk density remains uniformly low across almost the entire structure. When subdividing the region 100 subdivisions on the length and width dimensions and 75 subdivision on the height dimension, the density level distribution is

**Figure 3.**
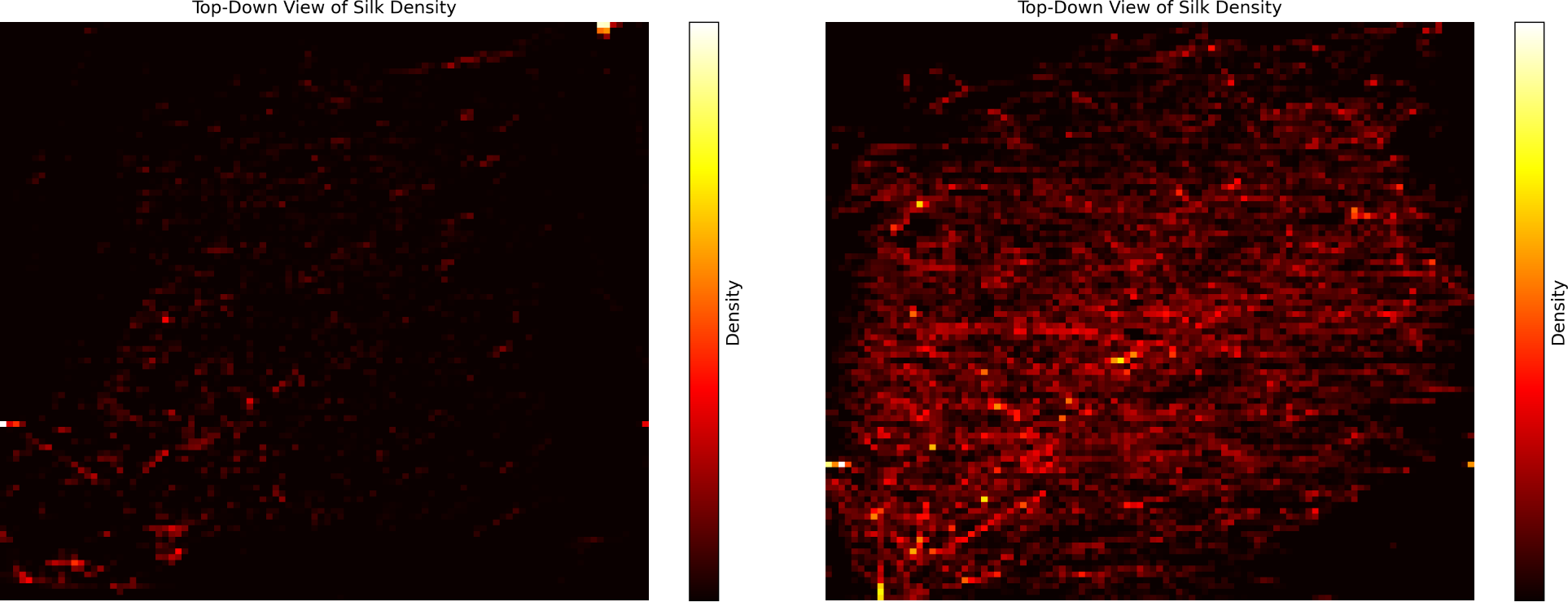
The figure on the left shows the silk density distribution for a web sample exhibiting a relatively low entropy value (0.5589 bits), calculated by using 10 quantization levels and excluding the empty level. The figure on the right shows the silk density distribution for a web sample exhibiting a relatively high entropy value (1.3300 bits), calculated by using 10 quantization levels and excluding the empty level.

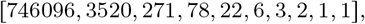

where the first entry represents the number of blocks with 0 density, the next entry represents the number of blocks with the lowest positive density level, and so on. This distribution indicates that most of the space is empty, with the lowest density level remaining dominant.

By contrast, the high-entropy web (Figure 3, right) shows greater heterogeneity. When subdividing the region 100 subdivisions on the length and width dimensions and 75 subdivision on the height dimension, the density level distribution is

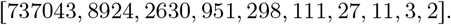

This distribution indicates that most of the space is still empty, with the lowest density level remaining relatively dominant; however, the spider employed a substantially wider range of density levels throughout the structure.

It should be noted that, in theory, if the majority of the web has high silk density, the entropy would still remain low due to consistency. However, our observations indicate that as the spider invests more energy in adding silk to the web, the additional silk density is distributed in a relatively random manner. This randomness prevents the density levels from concentrating on specific values, thereby increasing the entropy as the spider enhances the web’s density. Additionally, while high-density silk on the boundary provides structural integrity, the variability in silk density elsewhere reflects the spider’s adaptive energy investment [27]. For example, varying silk densities can optimize the web for prey capture, as areas with lower density allow prey to escape more easily [1].

### 2.2 Mathematical Framework for Harmonicity

#### 2.2.1 Motivation for Harmonicity

*If you know your neighbors, do you know yourself?* An important aspect in spider web complexity is the spatial correlations in silk density. By examining how the density of silk in one region of the web relates to the density in neighboring regions, we can infer patterns and interpolate the distribution of silk throughout the web. In particular, we want to infer a silk density of a subdivided box *B*_*i,j,k*_ based on densities of its neighboring subdivided boxes.

There are some nice properties that a discrete *harmonic* function (which will be defined in the following section) satisfy. For example, the value of a harmonic function at a point is determined as the average of the values of its neighbors. In particular, harmonic functions obey the maximum principle, which states that the highest values of the function appear at its boundary, an observed but typically unquantified trait of sheet webs [17, 1].

#### 2.2.2 Mathematical Framework for Harmonicity Computation

Let *f* be a function on a lattice **Z**^3^, the set of integer triples. We call *f harmonic* if the value of *f* at a point is equal to the average of *f* over neighboring points:

##### Definition 2.4

(Harmonic functions). A function *f* : **Z**^3^ → **R** is *harmonic* if *f* p**x**q is equal to

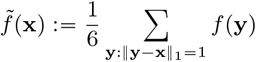

for any **x** ∈ **Z**^3^.

Equivalently, *f* is harmonic if and only if the *discrete Laplacian* Δ*f* vanishes identically [29, 30]:

##### Definition 2.5

(Laplacian and Harmonicity). For a function *f* : **Z**^3^ → **R**, the *Laplacian* of *f* is a function Δ*f* : **Z**^3^ → **R** defined as

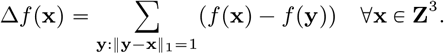

*Harmoncity* of *f* is defined as an *L*^2^-norm of Δ*f*.

As we mentioned above, we are interested in the spatial correlations in silk density. We want to infer the silk density of a box based on densities of its neighboring boxes. The simplest way to do this is by taking the average of neighbor densities, and this would work perfectly if and only if the silk density function *ρ* is *harmonic*.

By our earlier discussion, Δ*ρ* will be identically zero if and only if 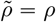. However, there is no reason for *ρ* to be a harmonic function a priori. Instead, we can use a generalized version of averages, and we consider *power means* as better alternatives. More precisely, we consider

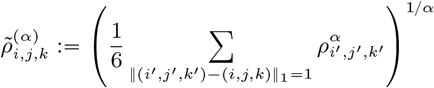

and see if we get a better inference results for some *α* ⩽ 0 other than *α =* 1, which corresponds to the usual mean. The *α*-power mean would give a zero error if and only if *ρ*^*α*^ is harmonic. This let us to define *α*-harmonicity of silk density:

##### Definition 2.6

(*α*-harmonicity of silk density). Define the *α-harmonicity* of a web *W* ‘s silk density *ρ* as ‖Δ(*ρ*^*α*^) ‖_2_, the harmonicity of *ρ*^*α*^.

We simply call 1-harmonicity as harmonicity. Like entropy, *α*-harmonicity depends on the choice of subdivisions *N* and the number of levels *L*. Under mild hypothesis, we can show that there always exists best *α* = *α* _*_ [0, ∞) that minimizes the *α*-harmonicity:

##### Theorem 2.7.

(Existence of the best *α*) If a web *W* with silk density satisfies the inequality

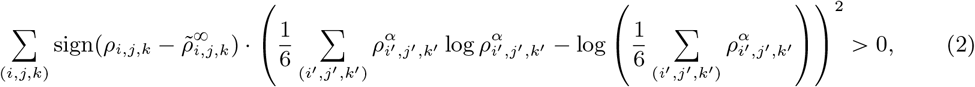

there exists finite *α= α*_*_ ⩽ 0 where *α*-harmonicity of silk density *ρ* is minimized.

Before we give a proof, we explain the condition (2). We found that the error function eventually increases for all of our webs, and this is a typical case in the following sense. The term after sign inside the summation in (2) is nonnegative by Jensen’s inequality. Also, sign 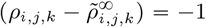 when 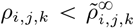, i.e. the silk density attains local maximum at *i, j, k*. In this case, all the neighboring indices (*i*^′^, *j*^′^, *k*^′^) with ‖ (*i, j, k*) (*i*^′^, *j*^′^, *k*^′^) ‖ _1_ 1 satisfy 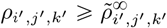 and the corresponding signs are plus. In other words, most of the signs in the summation are + 1, and this explains why the condition (2) is met.

*Proof*. First, the *α*-power average converges to the max function as *p* → ∞:

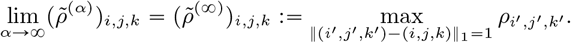

The logarithmic derivative of 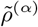 is

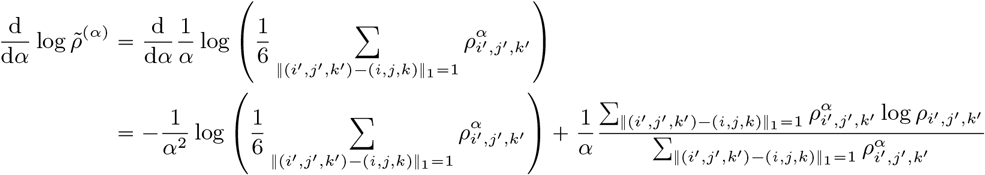

and by Jensen’s inequality applied to the convex function *f* (*x*) =*x* log *x* with 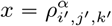, we have

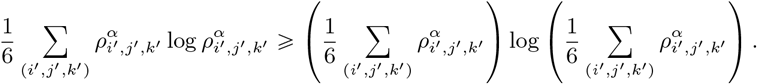

Hence the function 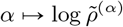 is increasing, as is 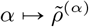.

Since 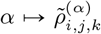 is increasing, 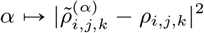 has two possible behaviors: increases for large *α* when 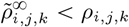, or decreases for large *α* when 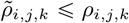. The derivative of *L*^2^-error 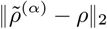 is

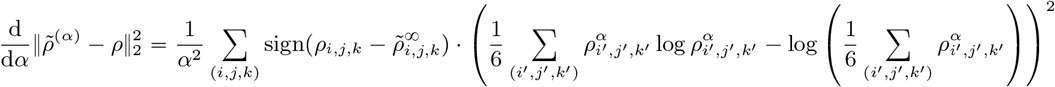

□

and the conclusion follows.

## 3 Methodology

### 3.1 Previous Methodologies

In [23], scanned images were stacked to form a 3D model of the web to understand the process of web construction. This process has been used to determine the purpose and importance of different stages in web construction, discern discrete stages of web construction, and estimate the mechanical properties of silk under stress in orb and tangle web species [31]. Instead of capturing and stacking individual frames sequentially as in [23] which takes 2 hours, we generate the entire scan as continuous video in 30 seconds and reconstruct the 3D volume in post-processing—drastically reducing acquisition time for each web scan.

Namazi used the fractal dimension and Shannon entropy to quantify the complexity and information content of orb weaving spiders to correlate cognitive function with web structure by interpreting the brightness level of digitized images [32]. However, web complexity was assessed using 2D projections of inherently three-dimensional structures, and important volumetric features such as out-of-plane geometries may be lost.

### 3.2 Specimen Collection and Web Construction

Local linyphiid species were used as test species for developing this new methodology and were hand-collected from the UC Berkeley campus. Not all specimens built webs during the allotted time, and only fully completed webs were included in the analysis. Collected spiders were kept in small containers for 1-2 weeks without food prior to introduction to experimental space to encourage web development [33]. The total number of spiders collected are 23, 5, and 13 for *N. digna, N*.*litigiosa*, and *M. dana*, respectively. Specimens were placed inside 10.16 × 10.16 × 10.16 cm (4” x 4” x 4”) acrylic boxes with clear sides, a black base, and no lid. Vaseline was coated on the inside top 2.54 cm (1”) of each side to prevent specimens from escaping. Box size was selected based the sizes of the species (all individuals measured less than 8mm in length excluding legs) and scanning capabilities of the experimental setup. Each spider was given at least two weeks in ambient conditions to complete a web, construction duration was not recorded in detail. Linyphiids construct webs over the course several days, completing full construction by seven days, and web completion was defined as the point at which no additional silk was added to the web after several consecutive days [33].

### 3.3 Web Scanning

Completed webs were placed into an enclosed experimental scanning apparatus blocking all outside light. Scanning was conducted by a laser (Quarton Inc. VLM-520-56) and camera (Canon EOS R5C mirrorless camera using a Canon RF 100mm F 2.8 Macro lens) mounted on a track at a fixed distance (0.67 meter). A small motor (Habow Technic Power-Functions XL-Motor-Set 8882) was used to advance the laser and camera simultaneously to scan the length of the experimental chamber from front to back, recording at 120 FPS for 20-30 seconds. An infrared laser distance sensor (DFROBOT DFRduin Uno v3.0[R3]) monitored the location of the camera relative to its start position to provide z-axis information for 3D reconstruction.

### 3.4 Data Processing and Extraction

Individual video frames representing two dimensional slices of the web were extracted for cleaning and analysis using a custom Python script [34]. For each video, a set cropped region was identified in each frame that excluded the box edges and background and accounted for any minute changes in elevation across the length of the box. Red and blue color values were reset to the minimum value and the image converted to grayscale. Background noise was removed using a minimum threshold filter of 0.55 max RGB value. Three-dimensional Point Cloud Data (PCD) representations of each web were created using the filtered data. Examples of the three-dimensional PCD plots are shown in Figure 7. All data are original scans collected as described above.

**Figure 4.**
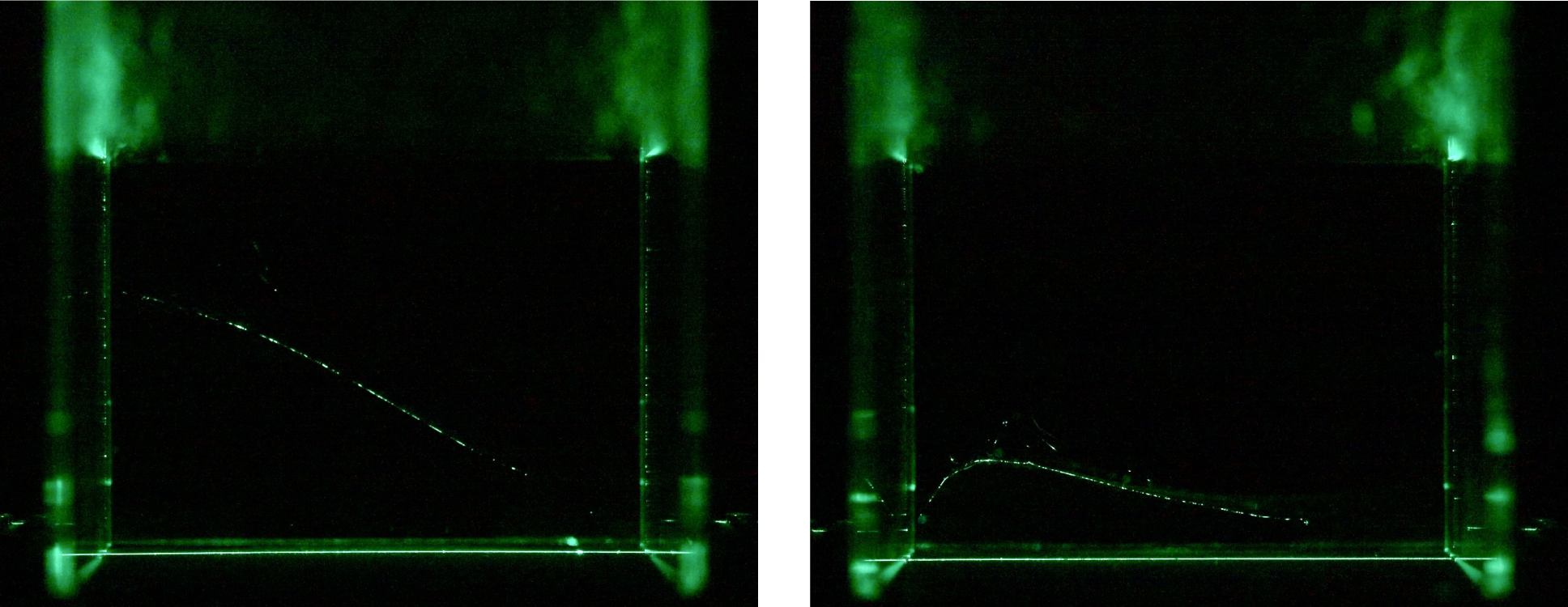
The figure on the left shows one representative frame of the web scan from the low entropy web in Figure 3 (left). The figure on the right shows one representative frame of the web scan from the high entropy web in Figure 3 (right).

**Figure 5.**
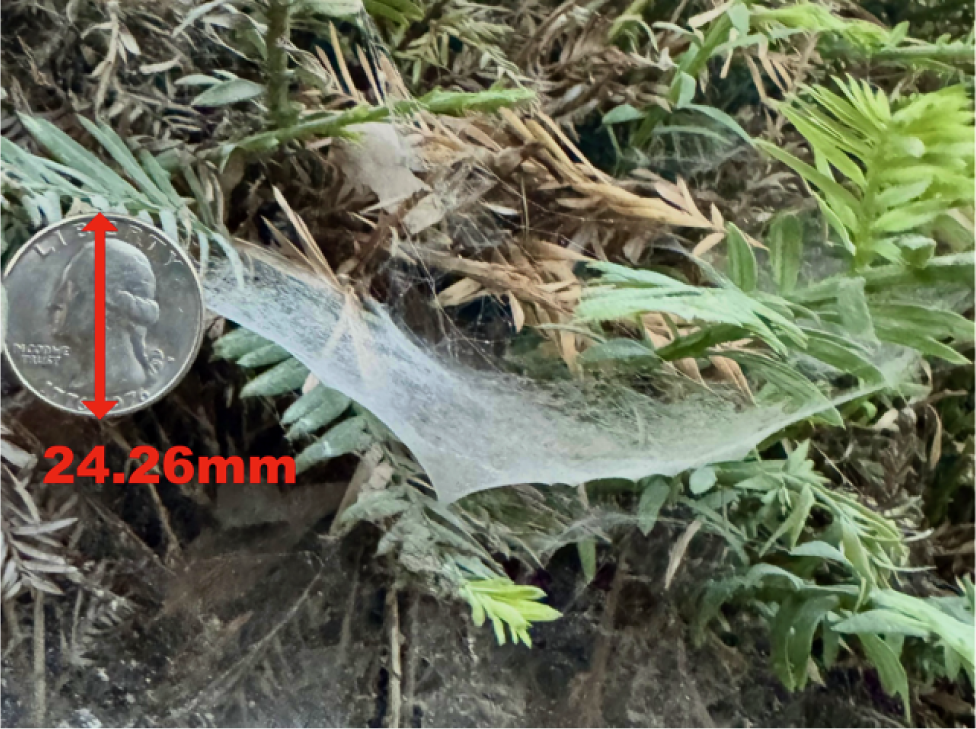
A linyphiid web on UC Berkeley campus (unidentified species). The web was dusted with cornstarch prior to photographing, quarter included for scale).

**Figure 6.**
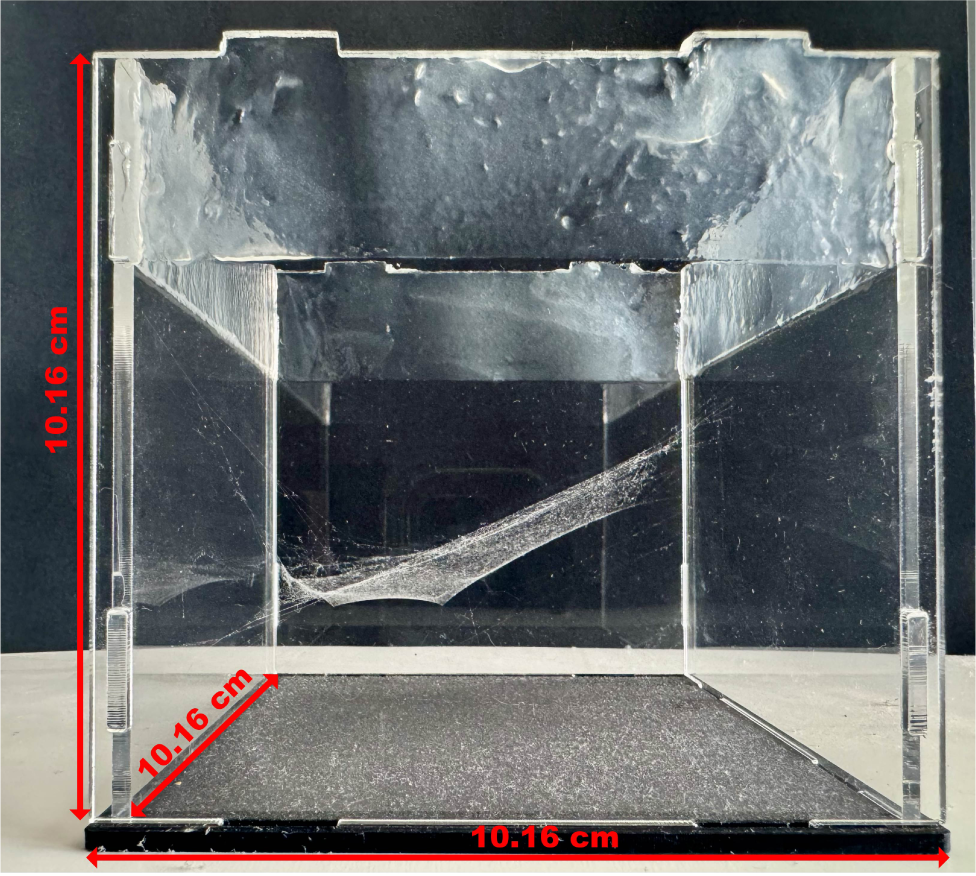
A representative image of a completed web in the experimental chamber. The web was dusted with cornstarch after scanning but prior to photographing for better visualization of the strucutre.

**Figure 7.**
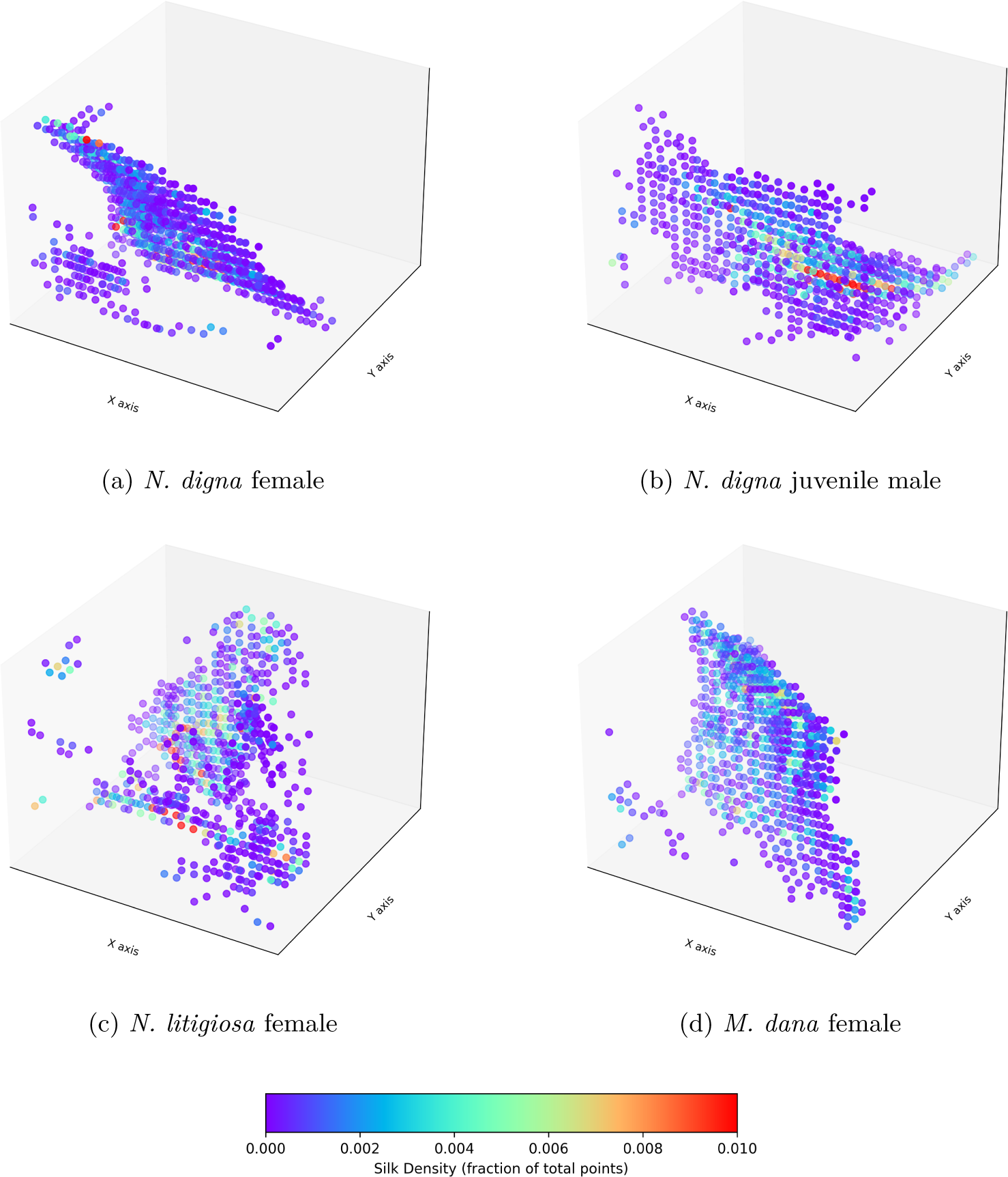
Visual representations of the silk density plots for four sheet webs from the three experimental species. These visualizations are derived from the PCD representation of the spider webs. The density in each region is computed based on the fraction of points within that region, relative to the total number of points. Purple colors represent less dense (or more sparse) areas of silk, while red areas indicate higher silk density.

**Figure 8.**
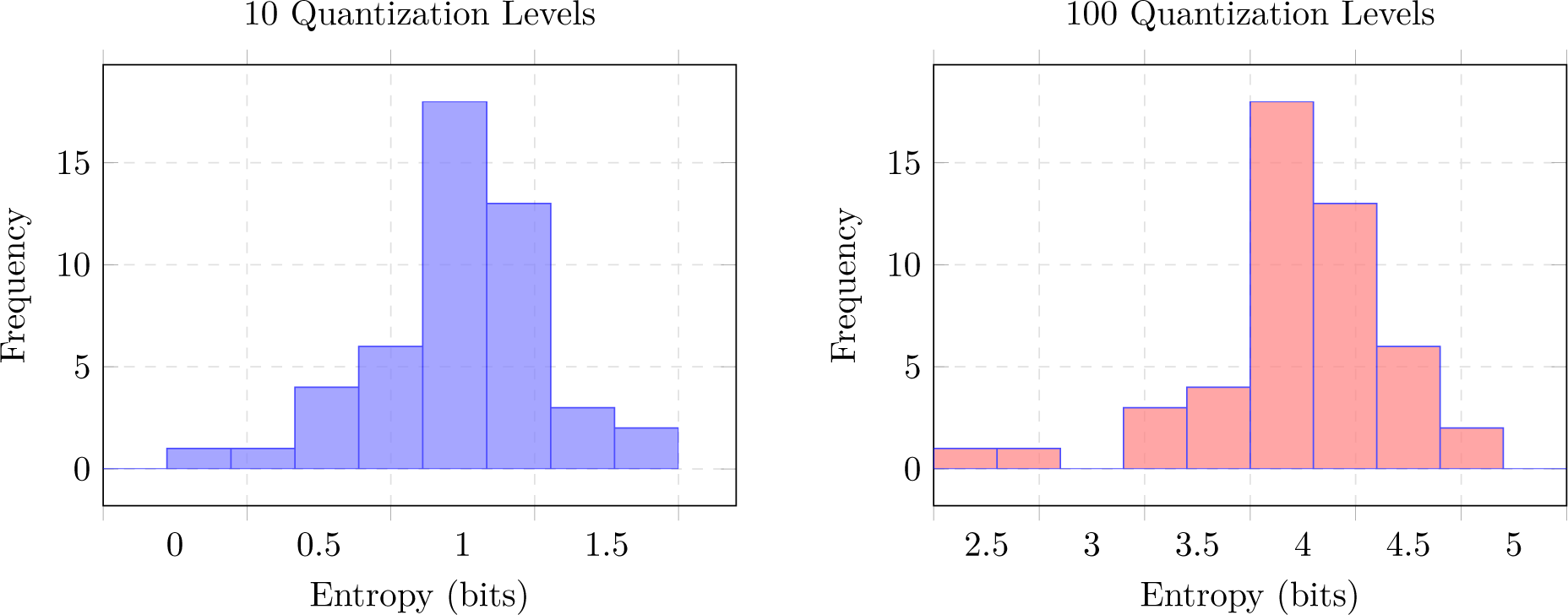
Histograms of entropy distributions computed with 10 and 100 quantization levels.

### 3.5 Quantification of Silk Density

We approximate the silk density in a region by counting the number of coordinate points in a subdivided box *B*_*i,j,k*_ (§2.1.2) of the corresponding Point Cloud Data document, divided by volume of *B*_*i,j,k*_. The historical method to determine silk density would be to physically cut each web over a partition and then measure the silk mass for each subdivision [35]. However, this approach poses practical challenges and destroys the web in the process. Using PCD representations yields us to gather accurate data while maintaining the web for further scanning or experimentation.

### 3.6 Statistical Analysis

The data for the entropy and harmonicity values for each web were analyzed in R 4.3 (http://R-project.org/) using the dplyr, ggplot2, lme4, and car packages: H (entropy), global harmonicity, and *α*_*_. Data exploration and analysis followed Zuur et al 2010 [36]. Outliers were removed based on IQR quartiles and the data was tested for normality using Shapiro–Wilk Normality Test (*p*-value = 0.441 for the entropy and *p*-value = 0.421 for harmonicity and *α*_*_) and Levene’s test for homogeneity of variance (for entropy: *F* (2, 38) = 0.364, *p* = 0.697; Harmonicity: *F* (2, 41) = 0.253, *p* = 0.778). Heteroscedasticity was tested using Breusch–Pagan test (*BP* = 9.095, *df* = 6, *p* = 0.168). We fit independent general linear mixed models to describe each response variable (H, global harmonicity, and *α*_*_) and assess the influence of species, genus, sex, and age, as well as the random effects of the date. The influence of each factor was modeled both independently and additively to test all possible combinations; statistical significance of the effect of the factors (species, genus, sex, age, and date) was not detected in any of the models. Comparisons between models were made using Akaike criterion, where the best fit model was identified based on the lowest AIC values. To assess multicollinearity, mixed models with multiple variables were tested using VIF and all GVIF^1/2(2·*f*)^ values were < 1.40 (species: 1.07; genus: 1.10; sex: 1.37; age: 1.40). Residual autocorrelation was checked with the Durbin–Watson test (*DW* = 2.43, *p* = 0.168).

## 4 Entropy of the Silk Density Results and Discussion

In this section, we address the question: *Is the silk density of a web consistent across its structure?* Our analysis reveals that the silk density exhibits a high degree of consistency, as indicated by the low entropy of the density level distribution - the average of the entropy *H* across the samples consistently converges to approximately 1.24 ± 0.22 bits (excluding zero-density-level) with respect to the number of samples for the spider species analyzed in this study, when using 10 quantization levels (Table 1). Based on our convergence analysis (Figure 1), which shows that the entropy reaches convergence once the number of subdivisions along the longest axes exceeds roughly 80, we therefore adopt 100 subdivisions (1 mm cell edge length) along length and width (and 75 along height) as a conservative yet efficient choice. To reduce sensitivity to the choice of subdivision size, we apply a moving average approach to the subdivision parameters. Specifically, the subdivisions are varied from 100 to 120 along the length and width dimensions, and from 75 to 100 along the height dimension. No significant effects of species (*F (*2, 38) = 1.897, *p =* 0.164), genus (*F*(1, 39) = 1.757, *p* 0.193), date (*F* (22, 18) = 1.111, *p* = 0.415), sex (*F*(2, 38) 0.756, *p* 0.476), and age (*F* (2, 38) = 2.600, *p =* 0.087) were identified. When the additive influence of species, date, sex, and age was modeled, no significant factors were identified (Best fit model - *F*(6, 36) 1.906, *p =* 0.1083). Further details on the specific sex and life stage for each specimen are included in Supplementary Table 3. One might wonder whether using more quantization levels than 10 could capture additional information about the silk density distribution. However, we computed entropy values up to 100 quantization levels, and the resulting distributions yielded a shape similar to that obtained with 10 levels. Increasing the number of density levels does not significantly alter the results under the current experimental conditions. The entropy value (*H*, excluding zerodensity-level) is 73 percent lower than the theoretical maximum of log_2_ 10 ≈ 3.32, reflecting an uneven distribution of density levels.

**Table 1:**
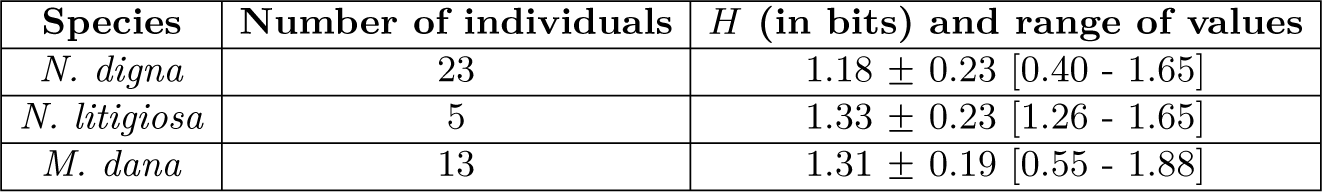
Calculated entropy values by species, presented as mean ± standard deviation [minimum value - maximum value]. For the computation of the entropy *H*, we remove empty space (level 0). Excluding level 0 allows us to compare the density levels only where silk is present, and more accurately reflects the organization of the sheet web.

The majority of the silk density falls within the lowest quantization level, suggesting an energy-saving behavior by the spiders. This low entropy may reflect a bias toward minimal silk usage, possibly prioritizing efficiency of resource use [20, 19, 37]. The production of silk and the construction of a web are energy intensive processes and are directly influenced by the physiological state of the spider and a number of ecological factors [6]. Among linyphiids species, silk density can range from quite sparse, as seen in *Laminacauda ansoni* to very dense in *Pocobletus versicolor*, where distinguishing individual thread lines is difficult [17]. Our results indicate that the sheet web spiders tested may optimize the amount of silk needed for effective prey capture while conserving energy; however, we cannot make decisive conclusions due to the limitations of the current study. The low silk density may be a factor of starvation, but this is unlikely, as starved spiders typically invest larger amounts of capture silk compared to sated individuals [6].

Alternatively, a higher entropy value would imply a more uniform use of different density levels, which could indicate that the spider may prioritize different factors of web performance across neighboring web sections related to varying functional purposes. For example, a dense region necessary for structural support adjacent to a sparse region of sticky silk for prey capture. This was not observed in the species used in this paper; however, the restrictions of the study may contribute to the observed consistency, as all specimens were kept in identical controlled conditions and had a relatively low sample size across all species, sexes, and life stages. Integrating greater numbers, species, and habitat diversity into future studies will either support low entropy as a universal feature of sheet webs or reject this conclusion and illuminate environmental and behavioral traits linked with differing entropy levels.

## 5 Harmonicity Results and Discussion

In this section, we address the question: *Can we infer silk density of a web by neighbors?* Using the mathematical framework described in Section 2.2, we calculated the global harmonicity of silk density (‖Δ ‖ _2_). No significant effects of species (*F* (2, 38) = 0.892, *p =* 0.419), genus (*F* (1, 39) = 1.757, *p* 0.193), date (*F*(22, 18) = 1.772, *p =* 0.111), sex (*F* (2, 38) = 0.6976, *p* 0.504), and age (*F* (2, 38) = 0.035, *p =* 0.965) were identified. When the additive influence of species, date, sex, and age was modeled, no significant factors were identified (Best fit model - *F*(6, 34) 0.576, *p* 0.746). For all webs, the average harmonicity value was 0.0039 ± 0.0017 (fraction of total points in PCD, Table 2). The harmonic behavior displayed in the silk density indicates that the silk is lain at similar densities across the surface of the sheet web, even when the specific shape of the web varies.

**Table 2:**
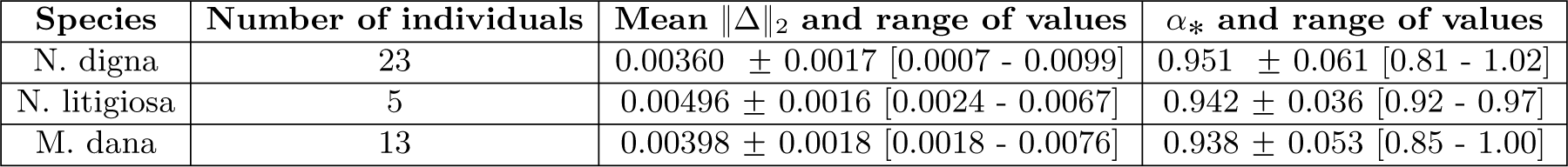
Harmonicity and *α* presented as mean ± standard deviation [minimum value - maximum value] for web scans separated by species. We use 20 subdivisions on the length and width dimensions and 15 subdivision on the height dimension. No significant differences were found between the species studied.

**Table 3:** Entropy values with 10 and 100 quantization levels for each web sample.

| Date of Scan | Species | H (in bits)<br>with 10<br>quantization levels | H (in bits)<br>with 100<br>quantization levels | $\rho_{\max}$<br>(fraction of total points) |
| --- | --- | --- | --- | --- |
| October 4 | N. digna Female | 0.7321 | 3.5442 | 0.002936 |
| October 6 | N. digna Imm. male | 1.2152 | 4.2575 | 0.001768 |
| October 8 | N. digna Female | 0.3962 | 2.7902 | 0.007339 |
| October 8 | N. digna Female | 0.6771 | 3.4151 | 0.005489 |
| October 17 | N. digna Female | 1.3843 | 4.4956 | 0.000590 |
| October 17 | N. digna Female | 0.8343 | 3.7659 | 0.001171 |
| October 19 | N. digna Female | 1.0561 | 4.1044 | 0.001741 |
| October 20 | N. digna Female | 0.9638 | 4.1044 | 0.001898 |
| October 26 | N. digna Female | 1.3831 | 4.5158 | 0.001915 |
| October 30 | N. litigiosa Female | 1.2605 | 4.1959 | 0.003066 |
| October 31 | N. digna Male | 0.7365 | 3.4904 | 0.002842 |
| November 2 | N. digna Female | 1.0041 | 3.9213 | 0.004980 |
| November 2 | N. digna Imm. Male | 1.6466 | 4.8236 | 0.000893 |
| November 5 | N. digna Female | 1.1862 | 4.2653 | 0.001245 |
| November 7 | M. dana Female | 1.1417 | 4.2489 | 0.002180 |
| November 8 | N. digna Female | 1.1358 | 4.1916 | 0.001783 |
| November 9 | M. dana Female | 1.5135 | 4.6413 | 0.002602 |
| November 12 | M. dana Female | 1.2542 | 4.3772 | 0.001217 |
| November 12 | M. dana Female | 1.8817 | 5.0831 | 0.000801 |
| November 12 | N. digna Female | 1.2572 | 4.3169 | 0.003028 |
| November 14 | N. litigiosa Female | 1.4815 | 4.6102 | 0.001532 |
| November 14 | M. dana Female | 1.1975 | 4.2305 | 0.001510 |
| November 14 | M. dana Female | 1.1184 | 4.1445 | 0.003016 |
| November 14 | N. digna Imm. Male | 1.2412 | 4.2306 | 0.001353 |
| November 16 | N. digna Female | 1.1021 | 4.1178 | 0.001396 |
| November 16 | M. dana Female | 1.1154 | 3.8777 | 0.002801 |
| November 17 | M. dana Female | 1.3573 | 4.4621 | 0.001023 |
| November 19 | M. dana Imm. Male | 0.5589 | 3.0589 | 0.006165 |
| November 21 | N. litigiosa Female | 1.3176 | 4.4031 | 0.001893 |
| November 25 | M. dana Female | 1.1582 | 4.2412 | 0.001284 |
| November 25 | M. dana Imm. Male | 1.4670 | 4.5988 | 0.001217 |
| November 25 | N. digna Imm. Male | 1.4693 | 4.5388 | 0.003416 |
| November 28 | N. digna Female | 1.2420 | 4.2934 | 0.001801 |
| November 28 | N. digna Imm. Male | 1.1231 | 4.0810 | 0.002882 |
| November 29 | M. dana Imm. Male | 1.8521 | 5.0588 | 0.000643 |
| December 1 | M. dana Imm. Female | 1.5872 | 4.7705 | 0.000673 |
| December 1 | N. digna Female | 1.0345 | 4.0807 | 0.001768 |
| December 1 | N. digna Imm. Female | 1.3856 | 4.5027 | 0.001159 |
| December 1 | N. litigiosa Female | 1.6531 | 4.7949 | 0.000975 |
| December 2 | N. digna Imm. Male | 1.1196 | 4.1265 | 0.002142 |
| December 4 | N. digna Imm. | 1.3714 | 4.4994 | 0.001616 |
| December 5 | M. dana Imm. Male | 1.3638 | 4.4815 | 0.000902 |
| December 5 | M. dana Imm. | 0.8912 | 3.9435 | 0.001582 |
| December 5 | N. digna Imm. Male | 1.3576 | 4.4665 | 0.001180 |
| December 5 | M. dana Imm. Male | 1.5067 | 4.6986 | 0.000896 |
| December 5 | M. dana Imm. Female | 1.2364 | 4.2909 | 0.002303 |
| December 9 | M. dana Imm. Female | 1.1792 | 4.1993 | 0.002849 |
| December 10 | N. digna Male | 1.3831 | 4.4186 | 0.002048 |

**Table 4:**
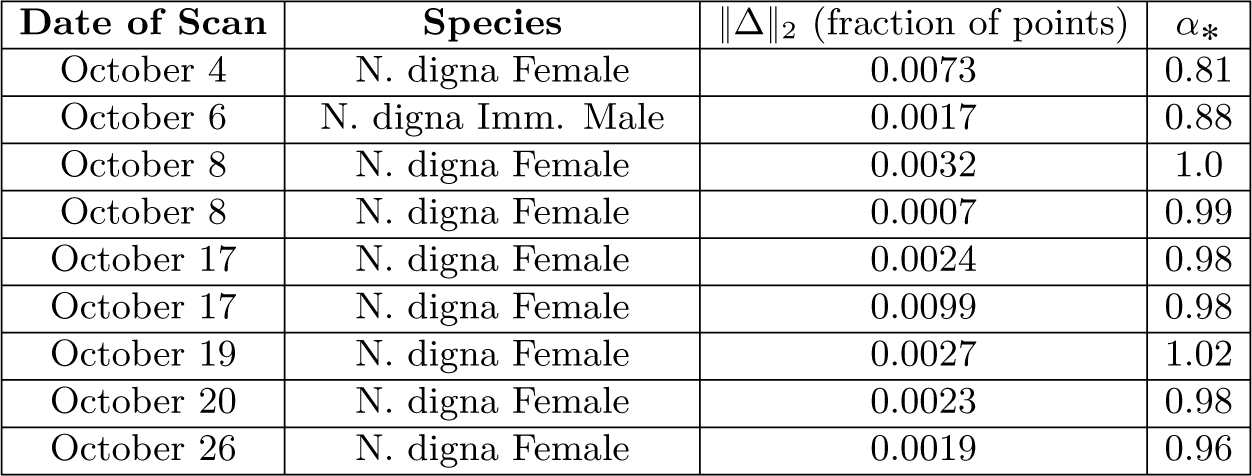
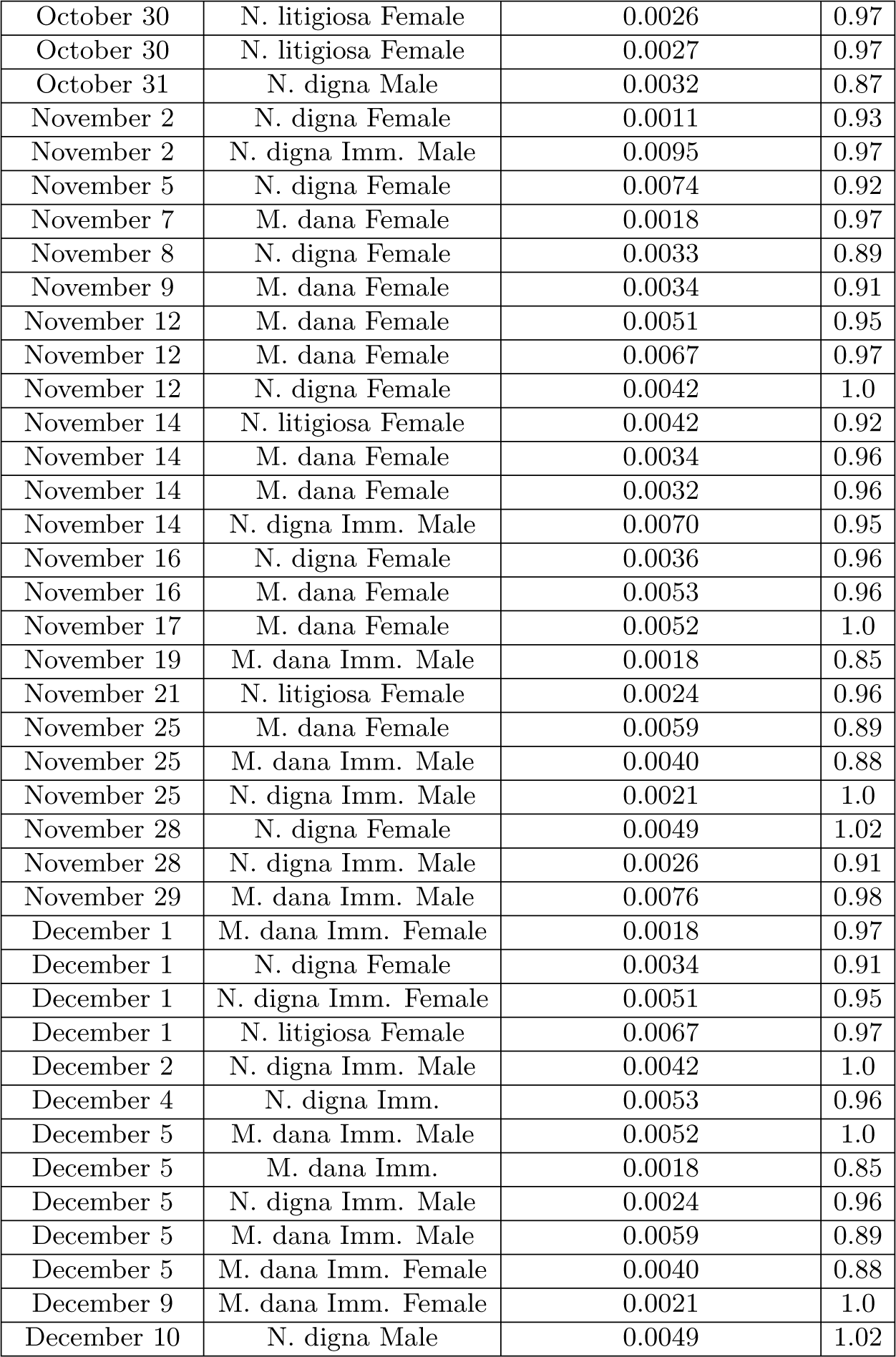
2-norm of the Laplacian for each web sample. We use 20 subdivisions on the length and width dimensions and 15 subdivision on the height dimension.

Also, we measure the *α*-harmonicity of each web and found the best *α = α*_*_ that minimizes the error, as mentioned at the end of Section 2.2 (see also Theorem 2.7). We compute the harmonicity of *ρ*^*α*^ by varying the *α* value in increments of 0.01 and find the *α* = *α*_*_ that minimizes ‖Δ(*ρ*^*α*^) ‖_2_.

We found that the average *α*_*_ is identically consistent to (‖Δ ‖_2_) over species, genus, sex, and age, at 0.95 ± 0.06 (Table 2). In other words, psilk densityq^0.95^ is approximately harmonic. *α*_*_ for the three linyphiid species.

## 6 Conclusions

In three species of linyphiid spiders, completed sheet-webs exhibit traits consistent with organization, optimization, and harmonicity, despite appearing random to the human eye. While the specifics of the organization and design are not yet known, the low entropy values provide evidence that the “unordered meshwork” is rather a deliberate network of silk fiber [7]. The limitations of the current study restrict our ability to draw broad biological and ecological conclusions, as only three, closely related species of Linyphiid were tested in a controlled space. Future studies integrating a wide diversity of species across varying environments will allow us to investigate how web structure relates to lifestyle, ecological conditions, and evolutionary history.

Our non-destructive approach allows us to draw quantifiable data from web structures. Moving forward, we can use these imaging and quantification methods to investigate web design in a wide variety of spider species across different environmental conditions and stressors. While only completed webs were scanned in this study, this non-destructive approach will allow us to obtain 3D scans of webs across every stage of construction, providing further insight into the step-by-step process used to create a viable model of the sheet-web. The 3D web scans can be analyzed in more detail to discern additional physical and geometric properties, and may be used to inform computational models of sheet-webs.

Most importantly, combining microscopic and macroscopic analysis of 3D web scans based on computed entropy and harmonicity values can bridge small-scale structural patterns with large-scale functional design that might reveal how spiders optimize their webs. From a microscopic perspective, metrics such as clustering coefficients, degree distributions, and modularity can be computed to analyze the graph structure of sheet webs. Viewing sheet webs as complex networks can provide deeper insights towards the specifics of thread organization and how this contributes towards the functionality of the web as both a shelter and a prey trap [38]. From a macroscopic perspective, efforts are already underway to derive equations that can accurately model the surfaces of the webs we have already scanned. With these equations, we will investigate whether sheet webs exhibit characteristics of minimal surfaces. We can also simulate these webs with such equations to test the viability of using webs as a model for applications in electronics and material science. With a sufficiently large dataset of spider web scans, machine learning models can be trained to generate web-like structures with similar properties.

Understanding the physical properties of sheet webs may also provide inspiration for engineering and architectural designs that minimize material usage while maintaining structural function. The ability to directly measure, experiment and manipulate webs during construction will open new doors into the study of spider behavior and ecology, and allow us to quantify what makes an effective web. Identifying the unseen patterns of nature is a powerful tool, and the non-destructive scanning technique and quantification of entropy and harmonicity described here have the potential to illuminate many undiscovered nuances in the structure and design of spider webs.

## Ethics

This work did not require ethical approval from a human subject or animal welfare committee.

## Author Contributions

Conceptualization: S. Lee, F. Lin, N. Sheu; Data Curation: P. Chang, J. Jiang, S. Lee, S. Li, F. Lin, K. Nagel, N. Sheu, G. Yang; Investigation: P. Chang, J. Jiang, S. Lee, S. Li, F. Lin, K. Nagel, N. Sheu, G. Yang; Methodology: J. Jiang, S. Lee, F. Lin, K. Nagel, N. Sheu, G. Yang; Software: P. Chang, J. Jiang, S. Lee, S. Li, F. Lin, K. Nagel, G. Yang; Formal analysis: J. Jiang, S. Lee, F. Lin, K. Nagel, N. Sheu, G. Yang; Validation: J. Jiang, S. Lee, F. Lin, K. Nagel, N. Sheu, G. Yang; Visualization: J. Jiang, S. Lee, F. Lin, K. Nagel, G. Yang; Writing-original draft: J. Jiang, S. Lee, F. Lin, K. Nagel, N. Sheu, G. Yang; Writing-review and editing: J. Jiang, S. Lee, F. Lin, K. Nagel, N. Sheu; Resources: J. Jiang, S. Lee, F. Lin, K. Nagel, N. Sheu, G. Yang; Supervision: F. Lin, N. Sheu; Project Administration: N. Sheu. All authors have read and agreed to the published version of the manuscript.

## Acknowledgements

We would like to thank Professor Per-Olof Persson and Professor Rosemary Gillespie for their valuable guidance and support. Also, the third author thanks Hyukpyo Hong for helpful discussions.

## 7 Supplementary Material

### 7.1 Tables of the Entropy Data

The complete table of entropy data is provided below. “Imm.” indicates immature spiders. If no sex is listed, it means the spider was too young to be sexed.

### 7.2 Table of the Harmonicity Data

The full table of harmonicity data is as follows.

## References

[1] Foelix R. Biology of spiders. Oxford university press; 2010.

[2] Römer L, Scheibel T. The elaborate structure of spider silk: structure and function of a natural high performance fiber. Prion. 2008;2(4):154–61.

[3] Ko FK, Wan LY. Engineering properties of spider silk. Handbook of properties of textile and technical fibres. 2018:185–220.

[4] Lewis RV. Spider silk: the unraveling of a mystery. Accounts of chemical research. 1992;25(9):392–8.

[5] Harmer AM, Blackledge TA, Madin JS, Herberstein ME. High-performance spider webs: integrating biomechanics, ecology and behaviour. Journal of the Royal Society Interface. 2011;8(57):457–71.

[6] Blackledge TA, Kuntner M, Agnarsson I. The form and function of spider orb webs: evolution from silk to ecosystems. In: Advances in insect physiology. vol. 41. Elsevier; 2011. p. 175–262.

[7] Benjamin SP, Düggelin M, Zschokke S. Fine structure of sheet-webs of Linyphia triangularis (Clerck) and Microlinyphia pusilla (Sundevall), with remarks on the presence of viscid silk. Acta Zoologica. 2002;83(1):49–59.

[8] Lott M, Poggetto VFD, Greco G, Pugno NM, Bosia F. Prey localization in spider orb webs using modal vibration analysis. Scientific Reports. 2022;12(1):19045.

[9] Hill PS, Lakes-Harlan R, Mazzoni V, Narins PM, Virant-Doberlet M, Wessel A, et al. Biotremology: studying vibrational behavior. vol. 6. Springer; 2019.

[10] Mortimer B. A spider’s vibration landscape: adaptations to promote vibrational information transfer in orb webs. Integrative and Comparative Biology. 2019;59(6):1636–45.

[11] Coddington JA. Cladistics and spider classification: Araneomorph phylogeny and the monophyly of orbweavers (Araneae: Araneomorphae, Orbiculariae). Acta Zoologica Fennica. 1991.

[12] Griswold CE, Coddington JA, Hormiga G, Scharff N. Phylogeny of the orb-web building spiders (Araneae, Orbiculariae: Deinopoidea, Araneoidea). Zoological Journal of the Linnean Society. 1998;123(1):1–99.

[13] Bond JE, Garrison NL, Hamilton CA, Godwin RL, Hedin M, Agnarsson I. Phylogenomics resolves a spider backbone phylogeny and rejects a prevailing paradigm for orb web evolution. Current Biology. 2014;24(15):1765–71.

[14] Kallal RJ, Kulkarni SS, Dimitrov D, Benavides LR, Arnedo MA, Giribet G, et al. Converging on the orb: denser taxon sampling elucidates spider phylogeny and new analytical methods support repeated evolution of the orb web. Cladistics. 2021;37(3):298–316.

[15] Dimitrov D, Lopardo L, Giribet G, Arnedo MA, Álvarez-Padilla F, Hormiga G. Tangled in a sparse spider web: single origin of orb weavers and their spinning work unravelled by denser taxonomic sampling. Proceedings of the Royal Society B: Biological Sciences. 2012;279(1732):1341–50.

[16] Catalog WS. World Spider Catalog Version 26; 2025. Accessed: 11 June 2025. Available from: https://wsc.nmbe.ch/.

[17] Hormiga G, Eberhard WG. Sheet webs of linyphioid spiders (araneae: linyphiidae, pimoidae): the light of diversity hidden under a linguistic basket. Bulletin of the Museum of Comparative Zoology. 2023;163(8):279–415.

[18] Peters HM, Kovoor J. The silk-producing system of Linyphia triangularis (Araneae, Linyphiidae) and some comparisons with Araneidae: structure, histochemistry and function. Zoomorphology. 1991;111:1–17.

[19] Schmitz A. Respiration in spiders (Araneae). Journal of Comparative Physiology B. 2016;186:403–15.

[20] Peakall DB, Witt PN. The energy budget of an orb web-building spider. Comp Biochem Physiol A. 1976;54(1976):187–90.

[21] Brown JH, Gillooly JF, Allen AP, Savage VM, West GB. Toward a metabolic theory of ecology. Ecology. 2004;85(7):1771–89.

[22] Brown JH, Burger JR, Hou C, Hall CA. The pace of life: metabolic energy, biological time, and life history. Integrative and Comparative Biology. 2022;62(5):1479–91.

[23] Su I, Narayanan N, Logrono MA, Guo K, Bisshop A, Mühlethaler R, et al. In situ three-dimensional spider web construction and mechanics. Proceedings of the National Academy of Sciences. 2021;118(33):e2101296118.

[24] Herberstein ME, Tso IM. Evaluation of formulae to estimate the capture area and mesh height of orb webs (Araneoidea, Araneae). The Journal of Arachnology. 2000;28(2):180–4.

[25] Corver A, Wilkerson N, Miller J, Gordus A. Distinct movement patterns generate stages of spider web building. Current Biology. 2021;31(22):4983–97.

[26] Venter C, Haddad CR, Codron D. A novel approach to determine the surface area of buckspoor spider webs and other irregular-shaped two-dimensional objects. MethodsX. 2022;9:101904.

[27] Tanaka K. Energetic cost of web construction and its effect on web relocation in the web-building spider Agelena limbata. Oecologia. 1989;81:459–64.

[28] Shannon CE. A mathematical theory of communication. The Bell system technical journal. 1948;27(3):379–423.

[29] Duffin R. Basic properties of discrete analytic functions. Duke Math J. 1956;23(1):335–63.

[30] Benjamini I, Lovász L. Harmonic and analytic functions on graphs. Journal of Geometry. 2003;76:3–15.

[31] Su I, Buehler MJ. Mesomechanics of a three-dimensional spider web. Journal of the Mechanics and Physics of Solids. 2020;144:104096.

[32] Namazi HR. The complexity based analysis of the correlation between spider’s brain signal and web. ARC Journal of Neuroscience. 2017;2(3):34–44.

[33] Benjamin SP, Zschokke S. Homology, behaviour and spider webs: web construction behaviour of Linyphia hortensis and L. triangularis (Araneae: Linyphiidae) and its evolutionary significance. Journal of Evolutionary Biology. 2004;17(1):120–30.

[34] Lab G. spiderweb; 2025. [Online; accessed 26-Jan-2025]. https://github.com/normansheu/spiderweb.

[35] Sensenig A, Agnarsson I, Blackledge T. Adult spiders use tougher silk: Ontogenetic changes in web architecture and silk biomechanics in the orb-weaver spider. Journal of Zoology. 2011;285(1):28–38.

[36] Zuur AF, Ieno EN, Elphick CS. A protocol for data exploration to avoid common statistical problems. Methods in ecology and evolution. 2010;1(1):3–14.

[37] Kawamoto TH, Machado FdA, Kaneto GE, Japyassu HF. Resting metabolic rates of two orbweb spiders: A first approach to evolutionary success of ecribellate spiders. Journal of Insect Physiology. 2011;57(3):427–32.

[38] Manoj B, Chakraborty A, Singh R. Complex networks: A networking and signal processing perspective. Pearson; 2018.

